# Emergence of binocular disparity selectivity through Hebbian learning

**DOI:** 10.1101/221283

**Authors:** Tushar Chauhan, Timothée Masquelier, Alexandre Montlibert, Benoit R. Cottereau

**Affiliations:** Université de Toulouse, Centre de Recherche Cerveau et Cognition, Toulouse, France; Centre National de la Recherche Scientifique, Toulouse, France

## Abstract

Neural selectivity in the early visual cortex strongly reflects the statistics of our environment (Barlow, 2001; Geisler, 2008). Although this has been described extensively in literature through various encoding hypotheses (Barlow and Földiák, 1989; Atick and Redlich, 1992; Olshausen and Field, 1996), an explanation as to how the cortex develops the structures to support these encoding schemes remains elusive. Here, using the more realistic example of binocular vision as opposed to monocular luminance-field images, we show how a simple Hebbian coincidence-detector is capable of accounting for the emergence of binocular, disparity selective, receptive fields. We propose a model based on spike-timing dependent plasticity (STDP) which not only converges to realistic single-cell and population characteristics, but also demonstrates how known biases in natural statistics may influence population encoding and downstream correlates of behaviour. Furthermore, we show that the receptive fields we obtain are closer in structure to electrophysiological data than those predicted by normative encoding schemes (Ringach, 2002). We also demonstrate the robustness of our model to the input dataset, noise at various processing stages, and internal parameter variation. Taken together, our modelling results suggest that Hebbian coincidence-detection is an important computational principle and could provide a biologically plausible mechanism for the emergence of selectivity to natural statistics in the early sensory cortex.

## 1. Introduction

Sensory stimuli are not only the inputs to, but also shape the very process of, neural computation (Hebbs, 1949; Barlow, 1961). A modern, more rigorous extension of this idea is the efficient coding theory (Barlow and Földiák, 1989; Atick and Redlich, 1992), which postulates that the computational aim of early sensory processing is to use the least possible resources (neurones, energy) to code the most informative features of the stimulus (information efficiency). A direct corollary to the efficient coding hypothesis is that if the inputs signals are coded efficiently, the statistical consistencies in the stimuli should then be reflected in the organisation and structure of the early cortex (Geisler, 2008). The largest body of work on the efficient coding principle lies within the visual sensory modality. In the context of vision, a number of studies have shown that information-theoretic constraints do indeed predict localised, oriented and band-pass representations, akin to those reported in the early visual cortex (Olshausen and Field, 1996). Over the years, several studies have shown how properties of natural images can not only explain neural selectivity (Olshausen and Field, 1996, 1997; Furmanski and Engel, 2000; Karklin and Lewicki, 2009; Samonds et al., 2012; Okazawa et al., 2015), but also predict behaviour (Webster and Mollon, 1997; Howe and Purves, 2002; Geisler, 2008; Geisler and Perry, 2009; Girshick et al., 2011; Cooper and Norcia, 2014; Burge and Geisler, 2015; Sebastian et al., 2017).

Nearly all these studies, however, rely on images of natural scenes acquired using a single camera – effectively using a 2-D projection of a 3-D scene. Although this representation captures the luminance-field statistics of the scene; however, it does not fully reflect how visual data is acquired by the human brain. Humans are binocular, and use two simultaneous retinal projections which enable the sensing of disparity (differences in retinal projections in the two eyes). Disparity, in turn, can be used to make inferences about the 3-D structure of the scene such as calculations of distance, depth and surface-slant. Despite critical results in the analysis of luminance-field statistics, only a handful of studies (Hoyer and Hyvärinen, 2000; Burge and Geisler, 2014; Hunter and Hibbard, 2015; Goncalves and Welchman, 2017) have attempted to analyse the relationship between binocular projections of natural scenes and the properties of binocular neurones in the early visual system. Furthermore, these studies investigated the relationship between natural scenes and cortical selectivity in terms of a global optimisation problem which is *solved* in the adult brain, leading to cortical structures which encode relevant statistics of the stimuli. However, the question as to how these encoding schemes might actually emerge in the early sensory cortex remains, as yet, unanswered (Stanley, 2013). Here, we show how simple coincidence detection based on spike-timing dependent plasticity (Bi and Poo, 1998; Caporale and Dan, 2008) (STDP) could offer a biologically plausible mechanism for arriving at neural population responses close to those reported in the early visual system. We endow a neural network with a Hebbian STDP rule, and find that an unsupervised exposure of this network to natural stereoscopic stimuli leads to a converged population which shows single-cell and population characteristics close to those reported in electrophysiological studies (Anzai et al., 1999; Durand et al., 2002; Prince et al., 2002b; Sprague et al., 2015). Moreover, the emergent receptive fields differ from those obtained by optimisation-based methods, and are more representative of those reported in literature (Ringach, 2002). Thus, taken together, our findings suggest that known rules of synaptic plasticity are sufficient to explain the relationship between biases reported in the early visual system and the statistics of natural stimuli. Furthermore, they also provide a compelling demonstration of how simple biological coincidence-detection (König et al., 1996; Brette, 2012; Masquelier, 2017) could explain the emergence of selectivity in early sensory and multi-modal neural populations, both during and after development.

## 2. Materials and Methods

### Datasets

To simulate binocular retinal input, we chose two datasets – the Hunter-Hibbard dataset and KITTI. The main results are reported for the Hunter-Hibbard database of stereo-images (Hunter and Hibbard, 2015), available at https://github.com/DavidWilliamHunter/Bivis under the MIT license which guarantees free usage and distribution. This database consists of 139 stereo-images of natural scenes relevant to everyday binocular experience, containing objects like vegetables, fruit, stones, shells, flowers, and vegetation. The database was captured using a realistic acquisition geometry close to the configuration of the human visual system while fixating straight ahead with zero elevation (Figure 1A). The distance between the two cameras was close to the human inter-ocular separation (65 mm), and the two cameras were focussed at realistic fixation-points from 50 cm to several metres away from the cameras. The lenses had a large field of view (20°), enabling them to capture binocular statistics across a sizeable part of the visual field. Figure 1A also shows a red-cyan anaglyph of one representative scene from the database, while Figure 1B shows the images captured by the left and right cameras. Due to the geometry of the cameras, there are subtle differences in the two images (disparity) which provide important information about the 3-D structure of the scene. The mimicking of realistic acquisition geometry is crucial for capturing disparity statistics which resemble those experienced by human observers. For comparison, we also report results using the KITTI database (available at http://www.cvlibs.net/datasets/kitti/) which uses parallel cameras – a geometry which does not correspond to the human visual system (see Figure 6C and related discussion for more details).

**Figure 1:**
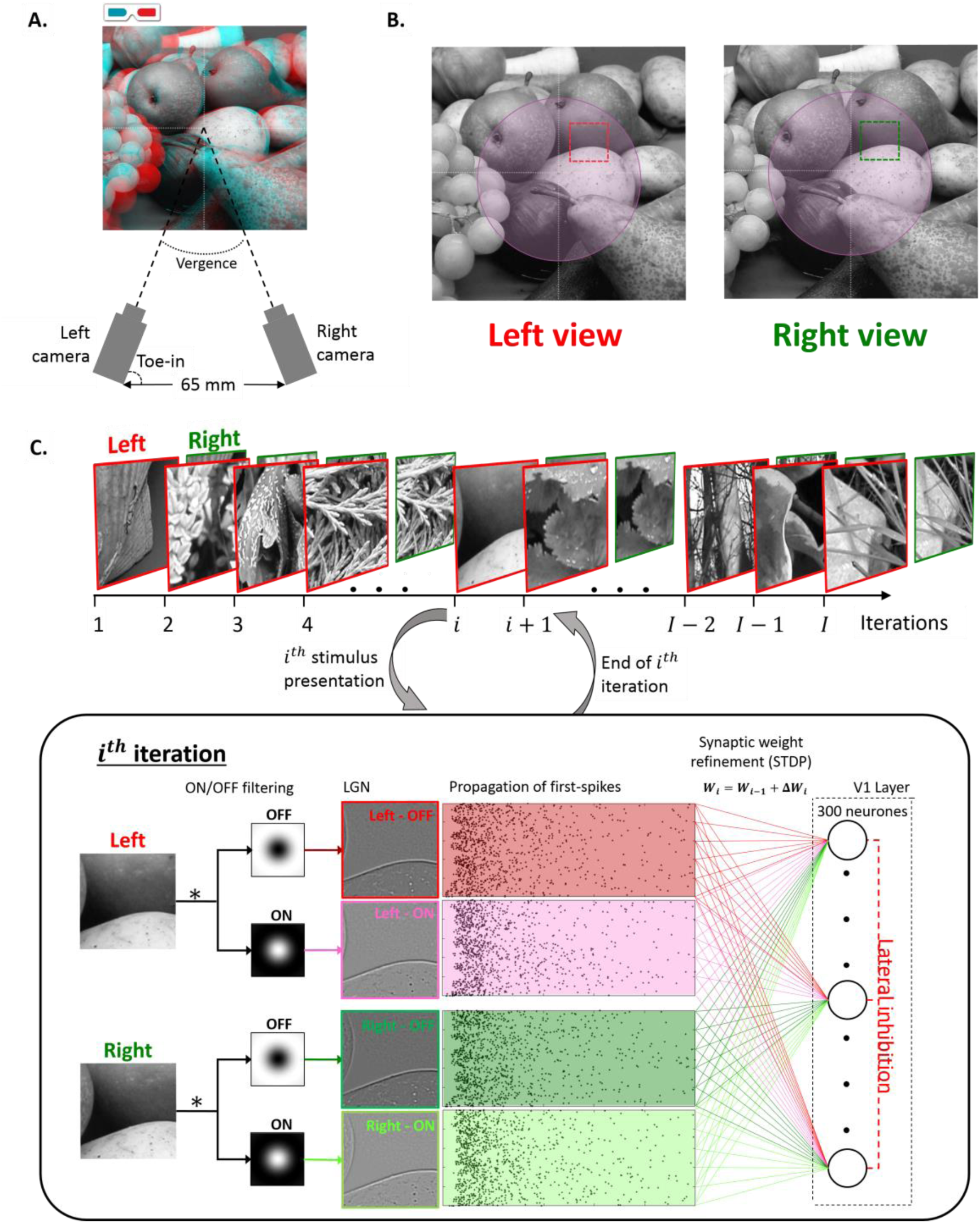
Processing pipeline. **A**. Acquisition geometry for the Hunter-Hibbard database. The set-up mimicked the typical geometry of the human visual system (see more details in the text). This geometry is crucial for capturing realistic disparity statistics from natural scenes. A stereoscopic reconstruction (red-cyan anaglyph) of a sample from the dataset is also shown. **B**. Input images captured by the left and the right camera for the scene shown in **A**. The subtle differences in the images (disparity) provide important cues about the 3-D structure of the scene. Depending on the simulation, a region-of-interest (ROI) in the visual field was identified, and inputs were sampled only within this ROI (foveal ROI illustrated in purple). Each sample consisted of two patches with equal retinal coordinates in the left and the right eye (illustrated in red and green respectively). All simulations presented in this paper were run using *N* = 100,000 samples each. **C**. The processing pipeline. The samples were processed sequentially, one per iteration. The processing of the *i^th^* sample is shown in a dashed box. First, the ON/OFF processing in the LGN was modelled using difference-of-Gaussian filters in the two eyes–leading to left-on, left-off, right-on and right-off maps. The activations of the four maps were then converted to first-spike relative latencies using a simple inverse operation. These first-spikes constituted the input to the V1 layer. The V1 layer consisted of 300 neurones connected to the LGN maps through Hebbian synapses endowed with spike-timing-dependent-plasticity (STDP). A lateral inhibition scheme was also implemented to ensure no more than one neurone fired per iteration. The first-spikes from the *i^th^* sample were propagated through the synapses, and this was followed by a change in synaptic weights dictated by the STDP algorithm (see text for details about the implemented STDP model). This concluded the processing of the *i^th^* sample, and was followed by the (*i* + 1)*^th^* sample, which was processed in a similar fashion.

### Input sampling

Random locations in the scenes were sampled to provide inputs to our model. In each simulation, the sampling was restricted to a specific region of interest (ROI). In this paper, we present results from four main ROIs – foveal (eccentricities less than 3°), peripheral (eccentricities larger than 6°), upper hemi-field, and lower hemi-field. Figure 1B shows the foveal ROI shaded in purple. Each sample consisted of two patches corresponding to the two eyes, both centred at the same random retinal coordinates. Figure 1B shows a random sample, with the left patch outlined in red and the right patch in green. The sizes of the patches varied with the ROI and can be interpreted as the initial dendritic receptive field of the V1 neurones, which was subsequently pruned by STDP. In the simulations presented in this paper, the foveal patches were 3° × 3° while the peripheral patches were 6° × 6°. 100,000 input samples were used in each simulation.

To simulate the fixation behaviour of typical strabismic amblyopes (Figure 3B), the left and right patches of each sample were no longer constrained to be centred at the same retinal coordinates, thereby introducing a misalignment between the two eyes (while still keeping within the ROI in each eye). The retinal correspondence between the ON and OFF LGN layers of the same eye was still maintained.

### Modelling of the retinogeniculate pathway

The computational pipeline employed by the model is shown in Figure 1C. The 100,000 samples were presented sequentially. The first stage of the model implemented the processing in the retinogeniculate pathway. The patches were convolved with ON and OFF centre-surround kernels to simulate the computations performed by the retinal ganglion cells (RGCs) and the lateral geniculate nucleus (LGN). The filters were implemented using Difference of Gaussian (DoG) kernels, and their receptive field sizes were chosen to reflect the size of representative LGN centre-surround magnocellular receptive fields (Solomon et al., 2002) – 0.3°/1° (centre/surround) for foveal simulations and 1°/2° (centre/surround) for peripheral simulations. The activity of each kernel can roughly be interpreted as a retinotopic contrast map of the sample. Four maps were calculated, corresponding to ON- and OFF-centre populations in the left and right monocular pathways. The activity of each sample was thresholded such that, on average, only a small portion (10%) of the LGN units fired. This activity was converted to relative first-spike latencies through a monotonically decreasing function (a token function *y* = 1/*x* was chosen, but all monotonically decreasing functions are equivalent (Masquelier and Thorpe, 2007)), thereby ensuring that the most active units fired first, while units with lower activity fired later, or not at all. Latency-based encoding of stimulus properties has been reported extensively in the early visual system (Celebrini et al., 1993; Gawne et al., 1996; Albrecht et al., 2002; Gollisch and Meister, 2008; Shriki et al., 2012), and allows for fast and efficient networks describing early visual selectivity (Thorpe et al., 2001; Masquelier and Thorpe, 2010; Masquelier, 2012). These trains of first-spikes (represented by their latencies) from the random samples constituted the input to the STDP network.

### V1: STDP neural network

The samples were presented to the network sequentially (Figure 1C). For each sample, the first-spikes from the LGN were propagated through plastic synapses to a V1 population of 300 integrate and fire neurones. The initial dendritic receptive fields of the neurones were three times the size of the LGN filters (foveal: 3° × 3°; peripheral: 6° × 6°). At the start, each neurone was fully connected to all LGN afferents within its receptive field through synapses with randomly assigned weights between 0 and 1. The weights were restricted between 0 and 1 throughout the simulation. The non-negative values of the weights reflect the fact that thalamic connections to V1 are predominantly excitatory in nature (Ferster and Lindström, 1983; Tanaka, 1985).

Figure 1C shows the processing for the *i^th^* sample in a dashed box. After the LGN spike propagation, the synaptic weights were updated using an unsupervised multiplicative STDP rule(Gütig et al., 2003). For a synapse connecting a pair of pre- and post-synaptic neurones, this rule is described by:

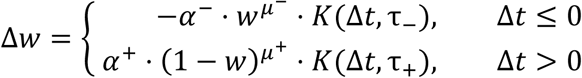

Here, a pre- and post-synaptic pair of spikes with a time difference Δ*t* introduce a change Δ*w* in the weight of the synapse which is given by the product of a temporal filter *K* and a power function of its current weight *w*. When a post-synaptic spike occurs shortly after a pre-synaptic spike (Δ*t* > 0), there is a strengthening of the synapse – also called long-term potentiation (LTP). Conversely, when the post-synaptic spike occurs before the pre-synaptic spike (Δ*t* ≤ 0), or in the absence of a pre-synaptic spike, there is a weakening of the synapse or a long-term depression (LTD). The LTP and LTD are driven by their respective learning rates *α*^+^ and *α*^−^. The learning rates are non-negative (*α*^±^ ≥ 0), and determine the maximum amount of change in synaptic weights when Δ*t* → ±0. The parameters *μ*^±^ ∈ [0,1] describe the degree of non-linearity in the LTP and LTD update rules. In practice, a non-linearity ensures that the final weights are graded and prevents convergence to bimodal distributions saturated at the upper and lower limits (0 and 1 in our case).

For the network used in the present study, the learning rates were fixed with *α*^+^ = 5 × 10^−3^ and *α*^+^/*α*^−^ = (4/3). The rate ratio *α*^+^/*α*^−^ is crucial to the stability of the network, and was chosen based on previous work demonstrating STDP based visual feature learning (Masquelier and Thorpe, 2007). Our simulations show that the results presented in this paper are robust for a large range of this parameter (see Figure 6B for more details). Furthermore, we used a high nonlinearity for the LTP process (*μ*^+^ = 0.65) to ensure that we were able to capture fading receptive fields through continuous weights, and used an almost additive LTD rule (*μ*^−^ = 0.05) to ensure pruning of continuously depressed synapses. In both LTP and LTD updates the weights were maintained in the range *w* ∈ [0,1]. Since our network relies only on first-spike latencies, the temporal filter *K* was approximated as a step-function (*τ*_±_ → ∞), making the update rule very efficient and fast. Furthermore, lateral inhibition through a winner-take-all scheme (Maass, 2000) was implemented such that, if any neurone fired during a certain iteration, it simultaneously prevented other neurones in the population from firing until the next sample was processed. This scheme leads to a sparse neural population where the probability of any two neurones learning the same feature is greatly reduced.

### Analysis of converged receptive fields

The post-convergence receptive fields of the STDP neurones were approximated by a linear summation of the afferent LGN receptive fields weighted by the converged synaptic weights. If a weight *w_ij_* connected the *i^th^* neurone to the *j^th^* LGN afferent with receptive field *ψ_j_*, the receptive field *ξ_i_* of the neurone was estimated by:

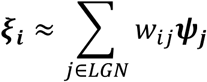

The receptive fields calculated using this method are, in principle, similar to point-wise estimates of the receptive field calculated by electrophysiologists. To better characterise the properties of these receptive fields, they were fitted by two dimensional Gabor functions. These Gabor functions are given by:

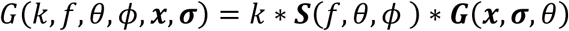

where, *k* is a measure of the sensitivity of the Gabor, ***S*** is a 2-D sinusoid propagating with frequency *f* and phase *ϕ* in the direction *θ*, and ***G*** is a 2-D Gaussian envelope centred at ***x***, with size parameter ***σ***, also oriented at an angle *θ*. A multi-start global search algorithm was used to estimate optimal parameters in each eye for each neurone. The process was expedited by using the frequency response of the cell (Fourier amplitude) to limit the search-space of the frequency parameter. For each neurone, the goodness of the Gabor fit was characterised using the maximum of the *R*^2^ goodness-of-fit values in the two eyes:

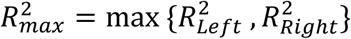

This ensured that the goodness-of-fit calculations were independent of whether the unit was monocular or binocular.

After the fitting procedure, the Gabor parameters were then used to characterise the structure and symmetry of the receptive field in the dominant eye (defined as the eye with the higher value of the Gabor sensitivity-parameter *k*). This was done by calculating the symmetry-invariant coordinates proposed by Ringach (2002). These coordinates (referred to as Ringach coordinates or RC from here on) are derived from the Gabor parameters by:

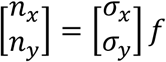

where, *σ_x_* is the spread of the Gaussian envelope along the direction of sinusoidal propagation, *σ_y_* is the Gaussian spread perpendicular to *σ_x_*, and *f* is the frequency of the sinusoid. While *n_x_* is a measure of the number of lobes in the receptive field, *n_y_* reflects the elongation of the receptive field perpendicular to the sinusoidal carrier. Typical cells reported in cat and monkey have both RC coordinates (*n_x_* and *n_y_*) less than 0.5 (Ringach, 2002).

The responses of the converged neurones to disparity (disparity tuning curves or DTCs) were characterised using two methodologies: binocular correlation (BC) of the left and right receptive fields, and estimation of responses to random dot stereograms (RDSs). In the BC method, we estimated the DTCs by cross-correlating the receptive fields in the left and the right eye along a given direction. In the RDS method, we estimated the DTCs by calculating responses of the converged neurones to random dot stereograms (RDSs) at various disparities. The RDS patterns used for testing were perfectly correlated, with a dot size of about 15′ visual angles, and a dot density of 0.24 (i.e., 24% of the stimulus was covered by dots, discounting any overlap). The DTC was calculated by averaging the response over a large number of stimulus presentations (*N* = 10,000 reported here).

The responses were further quantified by a Binocularity Interaction Index (Ohzawa and Freeman, 1986; Prince et al., 2002b) (BII) given by:

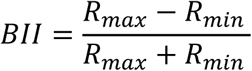

where *R_max_* and *R_min_* are maximum and minimum responses on the disparity tuning curve. BII values close to zero indicate that the neural response is weakly modulated by disparity, while higher values indicate a sharper tuning to specific disparity values. Furthermore, the phase and position disparity were estimated by methods commonly applied in electrophysiological studies – one-dimensional Gabor functions were fitted to the DTCs (for this estimation, we used the BC estimates as they are less subject to noise), and the phase of the sinusoid and the offset of the Gaussian envelope from zero-disparity were taken as estimates of the phase and position disparities respectively.

### Network convergence

The evolution of the synaptic weights with time was characterised by using a Convergence Index (CI). If Δ*w_ij_*(*t*) is the change in the weight of the synapse connecting the *i^th^* neurone to the *j^th^* LGN afferent at time *t*, the convergence index was defined as:

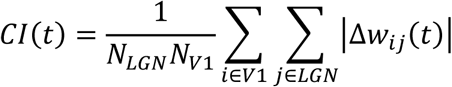

where *N_LGN_* and *N_V_*_1_ are the number of LGN units and the number of the V1 neurones respectively. The CI can be interpreted as a measure of the change in the synaptic weight distribution over time. Assuming a statistically stable input to a Hebbian network, the number of synapses undergoing a change in strength decreases with time. Thus, if our network is driven to convergence, CI should reduce asymptotically with time. To report the convergence for a single unit (say the *i^th^* V1 neurone), a modified form of the above index (*CI_i_*) was defined as:

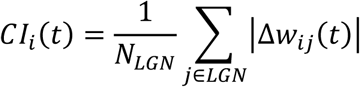

### Robustness analysis

Biological systems show a considerable resilience to factors such as noise and internal parameter variations (Burge and Jaini, 2017). We tested the robustness of our model using three approaches – 1) by introducing timing jitter at both the input and the LGN level, 2) by varying a key internal parameter of our network (the ratio of the LTP rate to the LTD rate), and 3) by comparing our results obtained using the Hunter-Hibbard dataset (realistic acquisition geometry) to results obtained using a dataset with non-realistic acquisition geometry (parallel cameras).

The robustness of the model to noise was tested by adding external (image) and internal (LGN) noise to the system. This simulated timing-jitter in the firing latencies from both the retina and the LGN. Gaussian noise with an SNR varying between -3dB (*σ_signal_* = 0.5 × *σ_noise_*) to 10dB (*_σsignal_* = 10 × *σ_noise_*) was used, and the performance of the network at various noise levels was characterised using two metrics – the convergence index (CI), and the 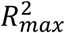 of Gabor fitting. While the CI can be used to evaluate the stability of the converged synaptic connectivity, the 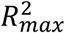 shows how well the receptive fields can be characterised by Gabor functions. Taken together, the CI and 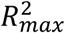 characterise the robustness of the network in terms of both the synaptic and the functional convergence.

The ratio of LTD to LTP rates (*λ* = *α*^−^/*α*^+^) is a critical parameter for the stability of the model as it determines the number of pruned synapses after convergence. If the probabilities of LTD and LTP events are *p*^−^ and *p*^+^ respectively, the learning rate *γ* can be approximated as

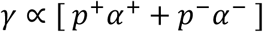

Given our initial simulation values of the LTD and LTP rates 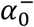 and 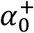 respectively, we defined 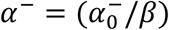 and 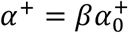, where the factor *β* was introduced to ensure that the two rates changed such that the overall learning rate *γ* remained stable. The effect of the LTP to LTD ratio on the convergence and functional stability of the network was then tested by running simulations for *λ* in logarithmic steps from 0.01 to 100 and estimating both the CI and 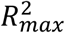 parameters. This simulated models where the LTD rates were from 1/100 to 100 times the LTP rates.

After testing the robustness of our model to noise and internal parameter variations, the aim was to test the model on a second dataset. As mentioned earlier, the Hunter-Hibbard dataset used in all our simulations was chosen because it has an acquisition geometry close to the human visual system. Thus, for this analysis we chose the KITTI dataset which was collected using a markedly different acquisition geometry. This dataset uses parallel cameras mounted 54 cm. apart, leading to a highly non-ecological, zero-vergence geometry. The aim of this analysis was to verify that the main results presented in this paper are not specific to the dataset, and that the model is capable of capturing natural disparity statistics whist being robust to acquisition geometry. Of course, in this case we did not expect the tunings of the converged neurones to match those reported in human electrophysiology (see Figure 6C and related discussion for more details).

### Decoding of disparity

The ability of the converged network to encode disparity was tested through a decoding paradigm. Random dot stereogram (RDS) stimuli at 11 equally spaced disparities between −1.5° and 1.5° were presented to the converged model, and the activity of the V1 layer was used to train, and subsequently test, linear and quadratic discriminant classifiers. We chose RDS stimuli because they do not contain any other information except disparity. A total of 10,000 stimuli per disparity were used, with a 70/30 training/testing split. 25-fold cross-validation testing was performed to ensure robust results. In this article, we report the detection probability (probability of correct identification) at each disparity. It must be noted that here, decoding through classifiers is only an illustrative representation of perceptual responses as it is based on inputs from very early visual processes and completely ignores important interactions such as cortico-cortical connections and feedback.

### Code accessibility

On acceptance of the manuscript for publication, the code will be freely made available on a public server (ModelDB and gitlab). During the review process, the editor and the reviewers will be given access to the code upon request.

## 3. Results

In this section, we present the results from simulations (10^5^ iterations per simulation) where binocular stimuli from specific ROIs were presented to our model. In the model, retinal inputs were filtered through an LGN layer, which was in turn connected to a V1 layer consisting of 300 integrate-and-fire neurones through STDP-driven synapses (see *Materials and Methods* for details). The main results are shown for the foveal ROI, while other ROIs (peripheral, upper-hemifield and lower-hemifield) are used to illustrate retinotopic biases.

### Foveal population

In Figure 2A, we present a representative unit from a foveally trained population of 300 V1 neurones. The first row of Figure 2A shows the initial receptive field of the neurone. Since the weights were set to random values, the resulting receptive field has no specificity in terms of structure or excitatory/inhibitory sub-fields. The second row of the figure shows the receptive field of the same neurone after STDP learning from 10^5^ samples (see also, Movie 1). The converged receptive field is local, bandpass, oriented. Clear on and off regions can be observed, and the receptive field closely resembles classical Gabor-like receptive fields reported for simple cells (Hubel and Wiesel, 1962; Jones and Palmer, 1987). Figure 2A also shows the Convergence Index (CI) of the neurone (bottom-left panel) which characterises the stability of its synaptic weights over the learning period. We find that the fluctuations in the average synaptic strength decrease over time to a very small value – indicating that the weight distribution has converged.

**Figure 2:**
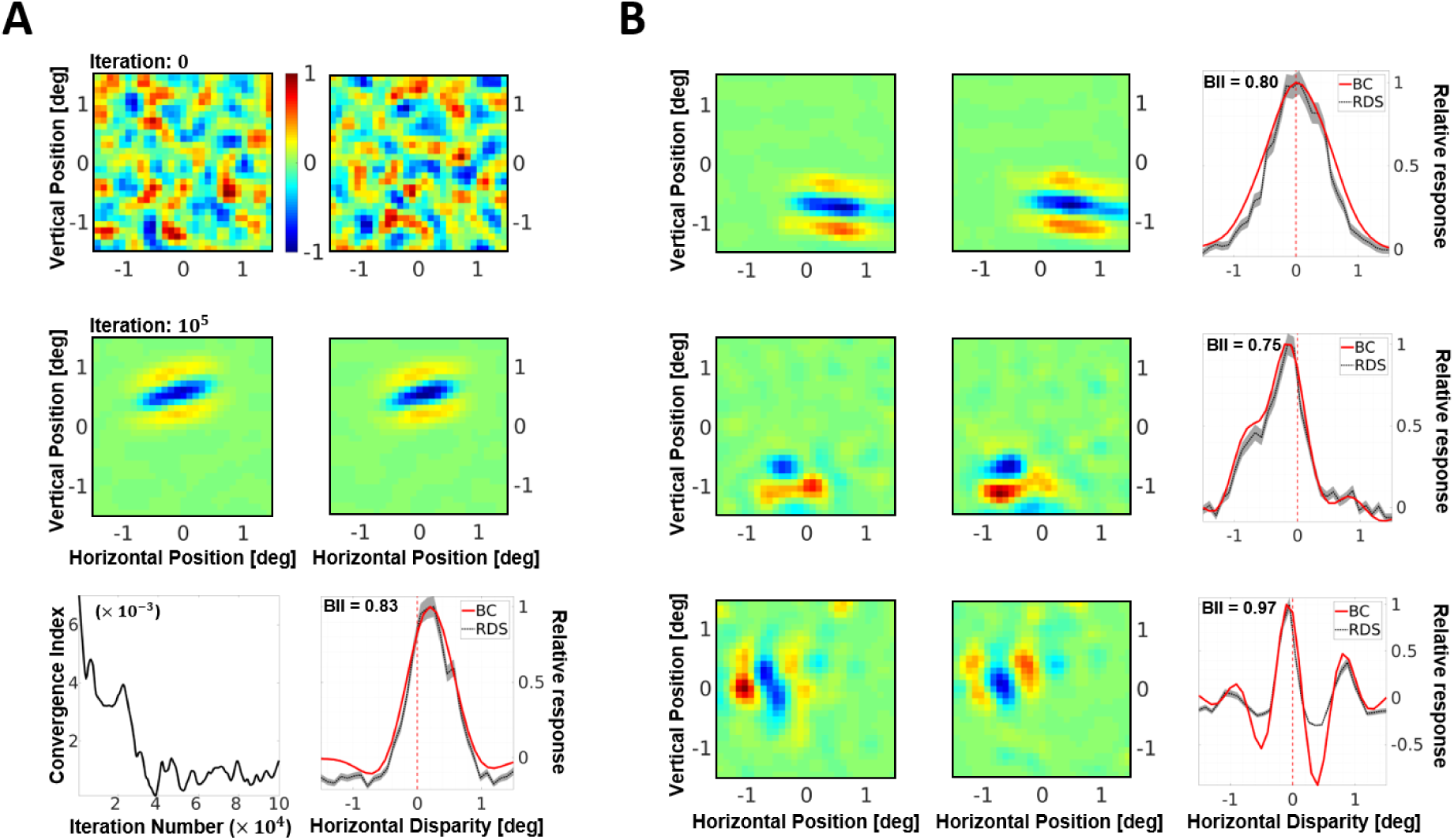
**A**. A representative neurone from the simulated population. Receptive fields in the left and the right eye before (first row), and after (second row) presentation of 10^5^ samples. The responses are scaled such that the maximum response is 1. The initial receptive fields are random, while the converged receptive fields show Gabor-like structure. The bottom-left panel shows the Convergence Index (CI), which measures the mean change in synaptic strength over time. Estimates of the Disparity tuning curve (DTC) for horizontal disparity are shown in the bottom-right panel. The neurone shown here is tuned to uncrossed (or positive) disparities. The DTC estimates using two techniques are presented –binocular correlation of the left and right receptive fields (red solid line), and averaging responses to multiple (*N* = 10^4^) presentations of random dot stereogram (RDS) stimuli (grey dotted line). The standard error for the RDS DTCs is shown as a grey envelope around the average responses. The Binocular Integration Index (BII) is also indicated within the graph. BII is a measure of the sensitivity of the neuronal response to disparity variations, with values close to 0 indicating low sensitivity, and values close to 1 indicating higher disparity sensitivity. See *Materials and Methods* for more details on the calculation of the receptive fields, DTCs and BII. **B**. Converged receptive fields and DTCs for neurones showing zero-tuned (first row), tuned near (second row) and phase-modulated (third row) responses to horizontal disparity.

In electrophysiology, a neurone with a binocular receptive field is often characterised in terms of its response to binocular disparity (Freeman and Ohzawa, 1992; Prince et al., 2002b, 2002a), i.e. its disparity tuning curve (DTC). The bottom-right panel of Figure 2A shows the DTC estimates for the selected neurone, estimated using both the BC (solid red line) and the RDS (dotted grey line, with standard error marked as a grey envelope) methods. The DTCs obtained from BC were very similar to those obtained through RDS stimuli. As the BC DTCs are, in general, less noisy, they were used to calculate all the disparity results presented in the article. This neurone shows a selectivity for positive or uncrossed disparities, indicating that it is more likely to respond to objects further away from the observer than fixation. It also has a high Binocularity Interaction Index (Ohzawa and Freeman, 1986; Prince et al., 2002b) (BII) of 0.83, indicating that there is a substantial modulation of its neuronal activity by binocular disparity (BII values close to 0 indicate no modulation of neuronal response by disparity, while values close to 1 indicate high sensitivity to disparity variations). Furthermore, Figure 2B shows three other examples of converged receptive fields and the corresponding DTCs (one neurone per row). The first neurone shows a tuning to zero-disparity (fixation), the second neurone to crossed or negative disparity (objects closer than the fixation), and the third neurone shows a tuning curve which is typically attributed to phase-tuned receptive fields. The receptive fields have a variety of orientations and sizes, and closely resemble two-dimensional Gabor functions.

### Receptive field properties

Since most converged receptive fields in the foveal simulation show a Gabor-like spatial structure, we investigated how well an ideal Gabor function would fit these receptive fields. Figure 3A shows the goodness-of-fit *R*^2^ parameter for the left and the right eye for all 300 neurones in the foveal population, with the colour code corresponding to the value 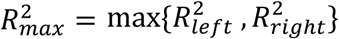. This value is maximal of the *R*^2^ parameter across the two eyes, and indicates how well the Gabor model fits the receptive field irrespective of whether it is monocular or binocular. A majority of the neurones lie along the diagonal in the upper right quadrant, showing an excellent fit in both eyes; with only few monocular neurones lying in the upper-left and bottom-right quadrants. It is also interesting to note that irrespective of the degree of fit, the *R*^2^ values for the left and the right eye are tightly correlated. This binocular correlation is non-trivial and cannot simply be attributed to learning from a stereoscopic dataset, as the inter-ocular correlations in the stimuli compete with intra-ocular correlations. To demonstrate this, in Figure 3B we show neurones from two simulated populations – the normal foveal population from Figure 3A (red dots), and a simulated amblyopic population (blue dots). Only the neurones with a reasonably good Gabor-fit 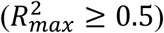 in each population are shown. The neurones in the amblyopic population occupy the top-left and bottom-right quadrants (indicating the neurones are monocular), as opposed to the binocular neurones from the normal population in the top-right quadrant. In the amblyopic population, the projections in the two eyes were sampled from different foveal locations within the same image (see *Materials and Methods*), thereby mimicking the oculomotor behaviour exhibited by strabismic amblyopes. Here, although neurones learn from left and right image-sets with the same widefield statistics, due to the lack of local inter-eye correlations they primarily develop monocular receptive fields – choosing features from either the left or the right eye.

**Figure 3:**
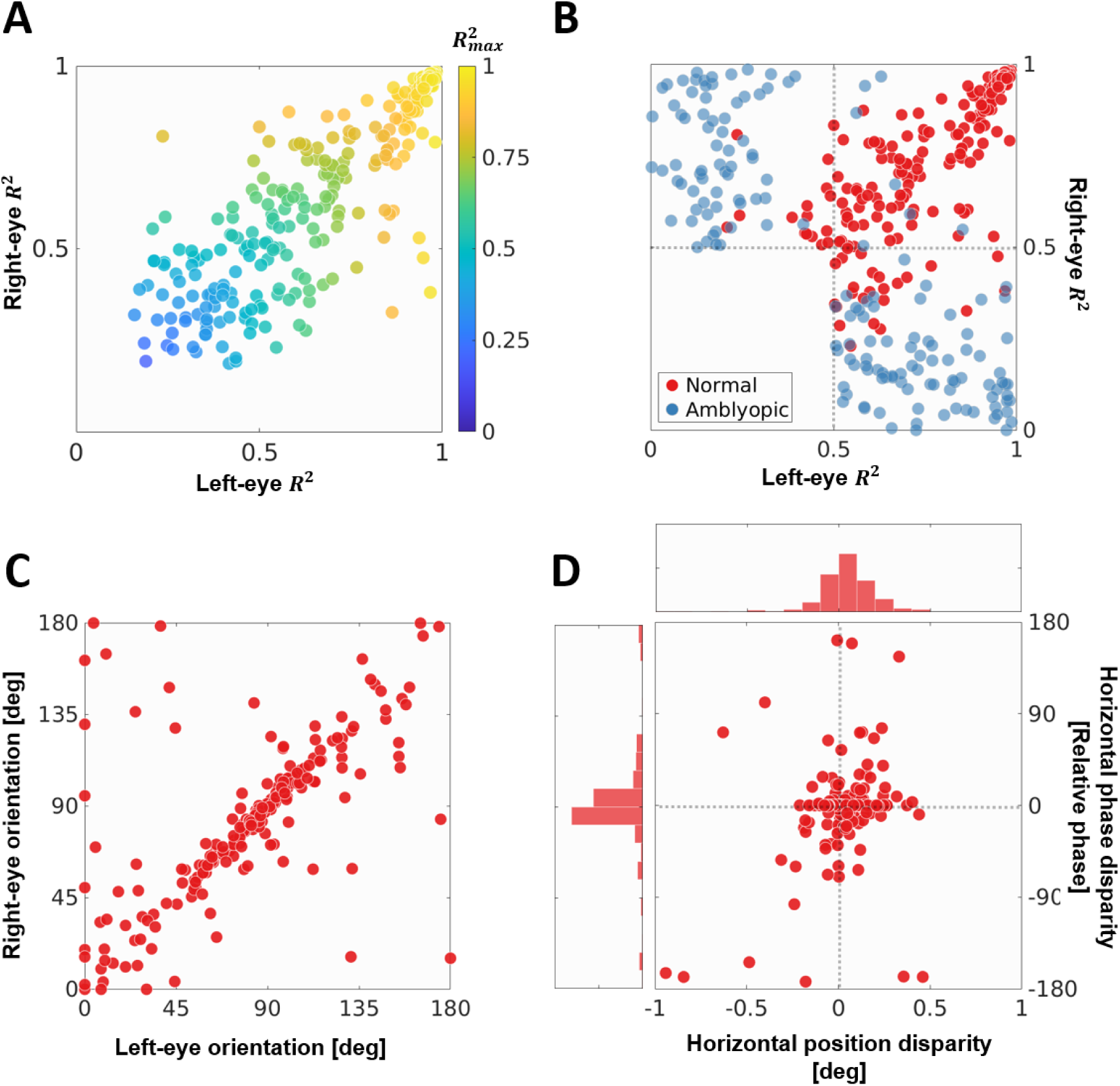
Characterising a simulated population of 300 foveal V1 neurones. **A**. The goodness-of-fit *R*^2^ criteria for Gabor-fitting in the left and the right eye. Each dot represents a neurone, while the filled colour indicates the quantity 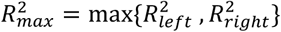, a measure of how well a Gabor function describes the receptive field irrespective of whether the neurone is monocular or binocular. **B**. Neurones 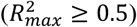 from normal (red dots) and amblyopic (blue dots) populations trained on foveal stimuli (see *Materials and Methods* for more details). **C**. The orientation parameter for the Gabors fitted to the left and the right eye receptive fields. Only neurones with binocular receptive fields defined by the criterion 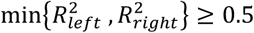 are shown. Note that the range of the angles is [0°, 180°], with 0° being the same as 180°. **D**. DTCs for the binocular neurones from **C**. were estimated by the binocular cross-correlation of the left and right-eye receptive fields under horizontal displacement. Gabors were fitted to the resulting curves, thereby giving a measure of the position and phase disparity in the receptive fields (Prince et al., 2002b, 2002a). This figure shows the position disparity plotted against the relative phase disparity. It also shows the histogrammes for the two types of disparities.

Figure 3C shows the receptive field orientations in the left and the right eye of binocular neurones in the foveal population. Here, we define a binocular unit by using the criterion: 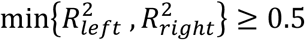, i.e., both eyes fit the Gabor-model with an *R*^2^ of at least 0.5. We see that there is a strong correspondence between the orientation in the two eyes (0° and 180° are equivalent orientations), with a large number of neurones being oriented along the cardinal directions, i.e., either horizontally or vertically. Though not shown here, a similar correspondence is found between the Gabor frequencies in the two eyes (the frequency range is limited between 0.75 and 1.25 cpd, due to the use of fixed LGN receptive field sizes), with an octave bandwidth of approximately 1.5 – a value close to those measured for simple-cells with similar frequency peaks (De Valois et al., 1982). Furthermore, Figure 3D shows the position and phase encoding of the horizontal disparities as derived by fitting Gabor functions to DTCs estimated by the BC method (see legend and *Materials and Methods* for details). Both position and phase are found to code for disparities between −0.5° and 0.5° visual angle (phase disparities were converted to absolute disparities using the frequency of the fitted Gabor), with most units coding for near-zero disparity.

In addition to edge-detecting Gabor-like receptive fields, more compact, blob-like neurones have also been reported in cat and monkey recordings (Jones and Palmer, 1987; Ringach, 2002). These receptive fields are not satisfactorily predicted by efficient approaches such as ICA and sparse coding (Ringach, 2002). In Figure 4, we present the distribution of Ringach-coordinates (RC, a symmetry-invariant descriptor for simple-cell receptive-fields, see *Materials and Methods* for details) for the dominant eye of our converged foveal population. The *n_x_* and *n_y_* axes approximately code for the number of lobes and the elongation of the receptive-field perpendicular to the periodic function (see *Materials and Methods* for more details). Most of the receptive fields in our converged population lie within 0.5 units on the two axes (blue shaded region) – two typical receptive fields in this region are shown (cyan markers). These receptive fields are not very elongated and typically contain 1-1.5 cycles of the periodic function. We also observe blob-like receptive fields (magenta markers) which display a larger invariance to orientation. In addition, we show three receptive fields which lie further away from the main cluster – along the *n_x_* axis with multiple lobes (brown markers), and along the *n_y_* axis with an elongated receptive field (purple marker). This RC distribution is close to those reported in cat and monkey recordings (Jones and Palmer, 1987; Ringach, 2002); while typically, RC distributions found by optimisation-based coding schemes such as ICA and sparse coding tend to lie to the right of this distribution (Ringach, 2002).

**Figure 4:**
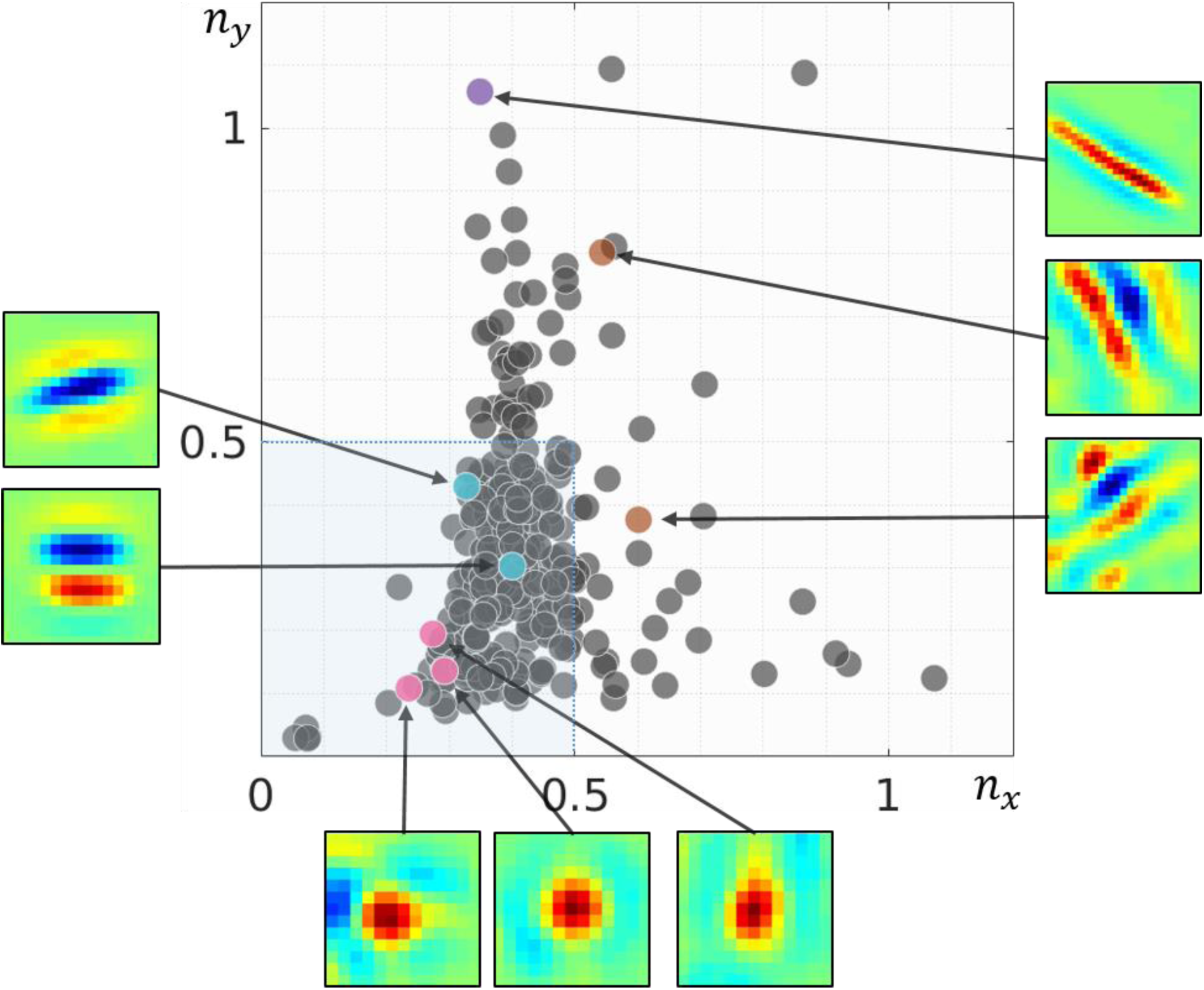
Ringach-coordinate (RC) distribution for the simulated foveal population (dominant eye). RC is a symmetry-invariant size descriptor for simple-cell receptive-fields (see *Materials and Methods* for a detailed description). High values along the *n_x_* axes convey large number of lobes in the receptive field, while high values along the *n_y_* axes describe elongated receptive fields. Receptive-fields for some representative neurones are also shown. The pink neurones are examples of blob-like receptive fields, while cyan markers illustrate 2-3 lobed neurones which constitute the majority of the converged population. Brown markers show neurones with a higher number of lobes, while the purple marker shows a neurone with an elongated receptive field. Most of the converged neurones lie within the square bounded by 0 and 0.5 along both axes (shaded in blue), something that is also observed in electrophysiological recordings in monkey and cat (Ringach, 2002).

### Biases in natural statistics and neural populations

The disparity statistics of any natural scene show certain characteristic biases which are often reflected in the neural encoding in the early visual pathway. In this section, we investigate whether such biases are also found in our neural populations. Figure 5A shows a histogram of the horizontal disparity in two simulated populations of neurones, one trained on inputs from the central field of vision (eccentricity ≤ 3°, shown in purple), and the other on inputs from the periphery (6° ≤ eccentricity ≤ 10°, shown in green). The peripheral population shows a clear broadening of its horizontal disparity tuning with respect to the central population (Wilcoxon rank-pair test: *p* = 0.046). As one traverses from the fovea to the periphery, the range of encoded horizontal disparities has indeed been shown to increase, both in V1 (Durand et al., 2007), and in natural scenes (Sprague et al., 2015).

**Figure 5:**
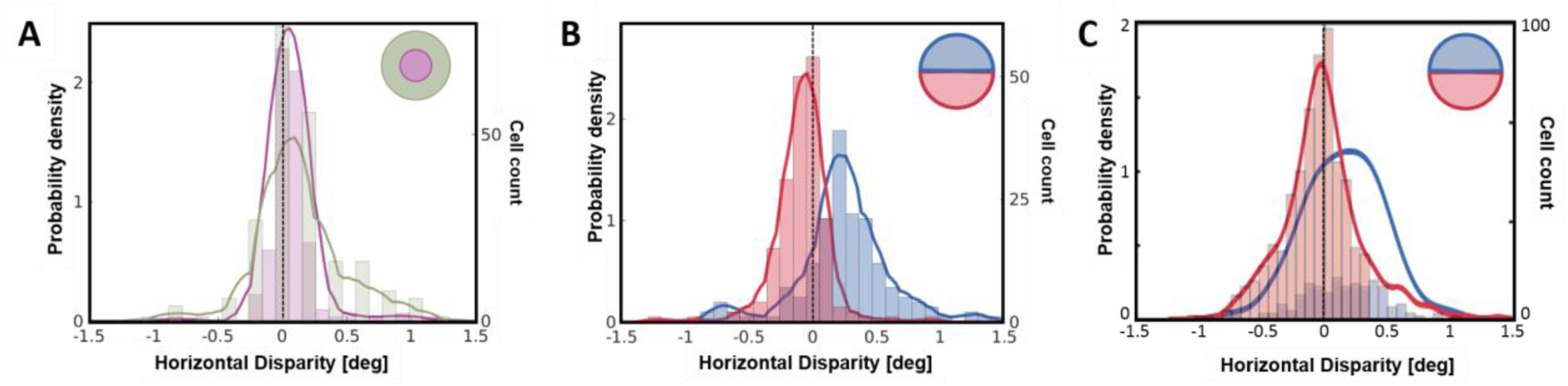
Retinotopic simulations and population biases in horizontal. **A**. Histograms and estimated probability density functions for horizontal disparity in two simulated populations of 300 V1 neurones each. The first population was trained on inputs sampled from the fovea (purple) while the second population was trained on inputs from the periphery (green). **B**. Histogram and probability density estimates of two populations of 300 neurones each, trained on inputs from the upper (blue) and lower (red) hemifield. **D**. A meta-analysis (Sprague et al., 2015) of results from electrophysiological recordings in monkey V1 neurones (820 neurones in total) by five separate groups. The neurones were split in two groups –neurones with upper-hemifield receptive fields are in blue, while those with receptive fields in the lower hemifield are in red.

Another statistical bias reported in retinal projections of natural scenes (Sprague et al., 2015) is that horizontal disparities in the lower hemifield are more likely to be crossed (negative) while the upper hemifield is more likely to contain uncrossed (positive) disparities. This bias is not altogether surprising considering that objects in the lower field of vision are likely to be closer to the observer than objects in the upper field (Hibbard and Bouzit, 2005; Sprague et al., 2015). A meta-analysis (Sprague et al., 2015) of 820 V1 neurones from five separate electrophysiological studies indicates that this bias could also be reflected in the binocular neurones in the early visual system. Figure 5C shows the results of this meta-analysis with the probability density for the neurones in the lower hemifield (in red) showing a bias towards crossed disparities while the upper hemifield (in blue) shows a bias towards uncrossed disparities. In comparison, Figure 5B presents histograms of the horizontal disparity in binocular neurones from two simulated populations – one trained on samples from the lower hemifield (red), and the other on the upper hemifield (blue). A comparison of Figure 5B and Figure 5C shows that both the range and the peak-tuning of the two simulated populations closely resemble those reported in electrophysiology. The peaks in Fig5B are at 0.17° for the upper hemifield and −0.02° for the lower hemifield (Wilcoxon rank-pair test: *p* = 2.15 × 10^−14^), while the meta-analysis by Sprague et al. (2015) shown in Fig5D reports the values as 0.14° and −0.09° respectively.

### Robustness to internal and external noise

In the results reported so far, the only source of noise are the camera sensors used to capture the stereoscopic dataset. To further test the robustness of our model we introduced additive white Gaussian noise ~*N* (0, *σ_noise_*) at two stages of the visual pipeline – external noise in the image, and internal noise in the LGN layer. The noise was manipulated such that the SNR varied in steps of approximately 3 dB from -3 dB (*σ_noise_* = 2 × *σ_signal_*) to 10 dB (*σ_noise_* = 0.1 × *σ_signal_*). Since the noise in these simulations induces timing jitter at the retinal and LGN stages, they are especially relevant for our model which relies on first-spike latencies. In Figure 6A we show the behaviour of the system under internal (in green colours) and external noise (red colours). The left and middle panels show the convergence index (CI) of the network, plotted against the iteration number (training duration) for the internal and external noise conditions respectively. The lightness of the colour indicates the level of noise (the darker the colour, the higher the noise). The right panel shows the mean 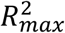, plotted against the SNR. We see that the CI approaches zero asymptotically in all conditions, indicating that the weights converge to a stable distribution for all levels of noise. The mean 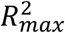, as expected, increases with the SNR (the higher the noise, the less Gabor-like the receptive fields). We see that the receptive fields converge to Gabor-like functions for a large range of internal and external noise, with the 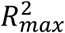 dropping below 0.5 only at SNR levels below 3 dB (*σ_noise_* ≥ 0.5 × *σ_signal_*).

**Figure 6:**
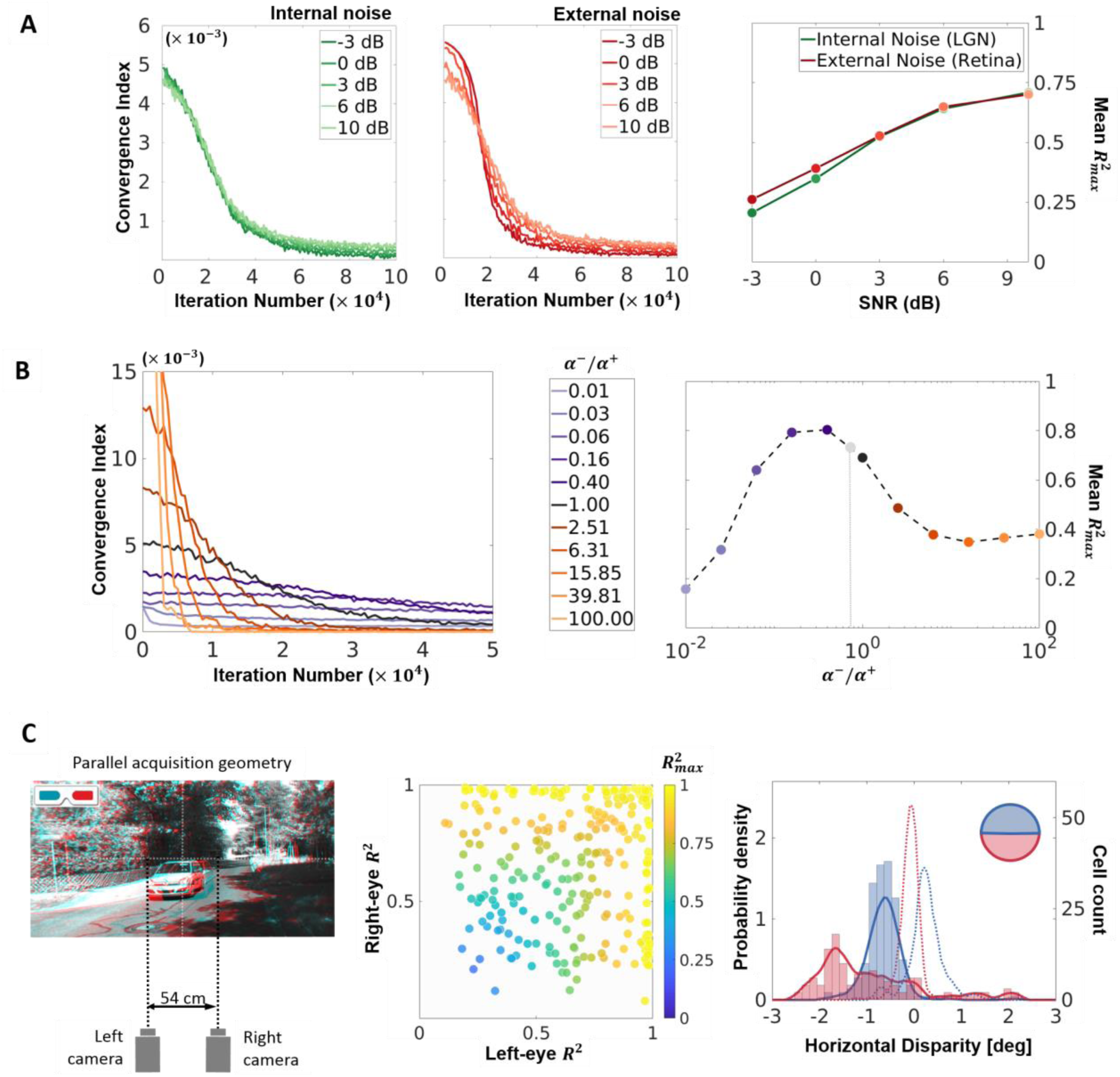
Robustness analysis. **A**. Internal and external noise. Additive white Gaussian noise (SNR levels between -3 and 10 dB) was introduced in the LGN activity (internal, in green colours), and the retinal input (external, in red colours). The left and middle panels show the Convergence Index (CI, see *Materials and Methods*) for the internal and external noise conditions respectively, plotted against the number of iterations (training period). In both graphs, the lightness of the curve codes for the SNR (the darker the curve, the higher the noise). For a stable synaptic convergence, the CI should approach zero asymptotically. The right panel shows the mean 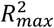 for the simulations (internal noise in green and external noise in red), plotted against the SNR. The higher the value of the mean 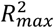, the closer the converged receptive fields are to Gabor-like receptive fields. **B**. Parameter variation. The ratio of the LTD rate (*α*^−^) to the LTP rate (*α*^+^), say *λ*, is a crucial parameter which determines the relative contributions of synaptic potentiation and depression to the overall learning rate. This parameter was varied logarithmically from 10^−2^ to 10^2^ (i.e., *α*^−^ = *α*^+^/100 to *α*^−^ = *α*^+^ × 100) to determine the robustness of the model to internal parameter variation. The left panel presents the CI for each level of *λ*, with purple colours coding for *α*^−^ ≤ *α*^+^ and orange/brown colours coding for *α*^−^ ≥ *α*^+^. The darker the colour, the closer the *α*^−^ to the *α*^+^, with black being *λ* = 1. The right panel shows the mean 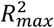 for the simulations plotted against *λ*, with the symbols following the same colour coding as the left panel. An additional grey dot is added to indicate the value of *λ* (=3/4) used for all the simulations presented in the paper (except this figure). The higher the mean 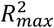, the more Gabor-like the converged receptive fields. **C**. A second dataset. The model was tested using a second dataset (KITTI, see *Materials and Methods* for details). The left panel shows the acquisition geometry of the KITTI dataset, and a cyan-red anaglyph reconstruction of a representative scene. The cameras were parallel and separated by 54 cm. –a geometry that does not resemble the human visual system. The middle panel shows the *R^2^* for the Gabor fit procedure on the left and right receptive fields of the converged neurones, with the colour of the symbol coding for the 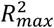 (compare to Figure 3A for the Hunter-Hibbard dataset). The right panel shows horizontal disparity tunings for two populations of neurones trained on the upper (blue) and lower (red) hemi-fields of the KITTI database. For comparison, the curves obtained using the Hunter-Hibbard database (Figure 5B) are plotted in dashed lines using the same colour scheme.

Overall, these analyses demonstrate that our approach is robust to a large range of noise induced timing-jitter at the input and LGN stages. While low noise conditions produce a synaptically converged network with Gabor-like receptive fields, high noise conditions produce a network which still converges, albeit to receptive fields which are no longer well-modelled by Gabor functions.

### Robustness to parameter variation

One of the key parameters of a Hebbian STDP model is the ratio (say, *λ*) of the LTD and LTP rates (*α*^−^ and *α*^+^ respectively). This parameter determines the balance between the weight changes induced by potentiation and depression, and thus directly influences the number of pruned synapses after convergence. The simulations presented in this article use *λ* = 3/4, a value based on STDP models previously developed in the team (Masquelier and Thorpe, 2007). In this section, we test the robustness of the model to variations in this important parameter. The value of *λ* was varied between 0.01 and 100 in log-steps (see *Materials and Methods* for the details). Figure 6B shows the results of these simulations. The left panel shows the CI, plotted against the iteration number (training period). The purple colours code for *α*^−^ ≤ *α*^+^ and the brown/orange colours code for *α*^−^ ≥ *α*^+^, with the intensity of the colour coding for the deviation from *λ* = 1. Once again, the synaptic weights converge for all values of the parameter, with convergence being quicker for higher values of *λ*. The right panel shows the mean 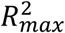 plotted against *λ*. The same colour code as the left panel is used, and an additional grey symbol indicates the performance of the model at *λ* = 3/4. The model produces Gabor-like receptive fields for a very broad range of *λ* values (for 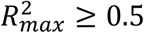 this range is roughly between 0.05 and 2), with the performance deteriorating for small values of *λ* (*α*^−^ is very low compared to *α*^+^) and saturating for *α*^−^ ≥ 6*α*^+^. This analysis shows that our model shows stable synaptic convergence for all tested values of *λ* (Figure 6B, left panel), and a functional convergence to Gabor-like receptive fields for a large range of the parameter (Figure 6B, right panel).

### Robustness to dataset acquisition geometry

All our simulations (except this section) use the Hunter-Hibbard database which was collected using an acquisition geometry close to the human visual system. The emergent receptive fields are disparity-sensitive, and closely resemble those reported in the early visual system. In this analysis, we prove that this emergence is not only a property of the dataset, but also of the human-like acquisition geometry. To do this, the model was tested on another dataset (KITTI, available at http://www.cvlibs.net/datasets/kitti/) which was collected using two widely-spaced (54 cm apart), parallel cameras – an acquisition-geometry which does not resemble the human visual system at all (left panel, Figure 6C). In the middle and right panel of Figure 6C, we present two results from this series of simulations. The middle panel shows the *R*^2^ of Gabor-fitting in the two eyes (symbols colour coded by 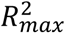) for a population trained on the foveal region of the KITTI dataset. We observe both monocular (high *R*^2^ in one eye, top-left or bottom-right quadrants) and binocular (high *R*^2^ in both eyes, top-right quadrant) neurones. In comparison, the converged receptive fields for the Hunter-Hibbard dataset are mostly binocular (Figure 3A). This is not surprising as the large inter-camera distance (54 cm as compared to 6.5 cm for the Hunter-Hibbard dataset) combined with the parallel geometry results in less overlap in the left and right images of the KITTI dataset, which further leads to weaker binocular correlations.

The right panel shows population disparity tunings for two separate populations trained on the upper (blue) and lower (red) hemi-fields of the KITTI dataset. As expected, the lower-hemifield population is tuned to disparities which are more crossed (negative) than the upper-hemifield population (see Figure 3C and the section *Biases in natural statistics and neural populations*). For comparison, the curves for the Hunter-Hibbard dataset (from Figure 3C) are plotted as dotted lines. The parallel geometry of the cameras in the KITTI dataset results in predominantly crossed disparities (the fixation point is at infinity, and hence all disparities are, by definition, crossed). Thus, while the upper vs. lower hemifield bias (see section *Biases in natural statistics and neural populations*) still emerges from our model, the population tunings no longer correspond to those reported in electrophysiological recordings (Sprague et al., 2015) (see Figure 3D).

### Decoding disparity through V1 responses

Recently, the effect of natural 3D statistics on behaviour has been modelled using labelled datasets in conjunction with task-optimised filters (Burge and Geisler, 2014) and feedforward convolutional networks (Goncalves and Welchman, 2017). Typically, such models are evaluated by how well they perform at classifying/detecting previously unseen data. In Figure 7A, we show the result (detection probability) of training two very simple classifiers on the activity of the converged V1 layer from our model, using RDS stimuli at 11 equally spaced disparities between −1.5° and 1.5° (see *Materials and Methods* for details). RDS stimuli were used because they contain no other meaningful information except disparity. The first, a decoder based on linear discriminant analysis (green), performs well above chance (dashed red), especially for realistic values of disparity between −0.5° and 0.5°. Interestingly, when the complexity of the decoder is increased slightly by including quadratic terms (blue), one observes a substantial increase in discrimination performance. This is not surprising as our neurones are linear units similar to simple cells, and a non-linear processing of their activity makes the decoding units conceptually closer to sharply tuned complex cells (Ohzawa et al., 1997; Cumming and DeAngelis, 2001; Henriksen et al., 2016).

**Figure 7:**
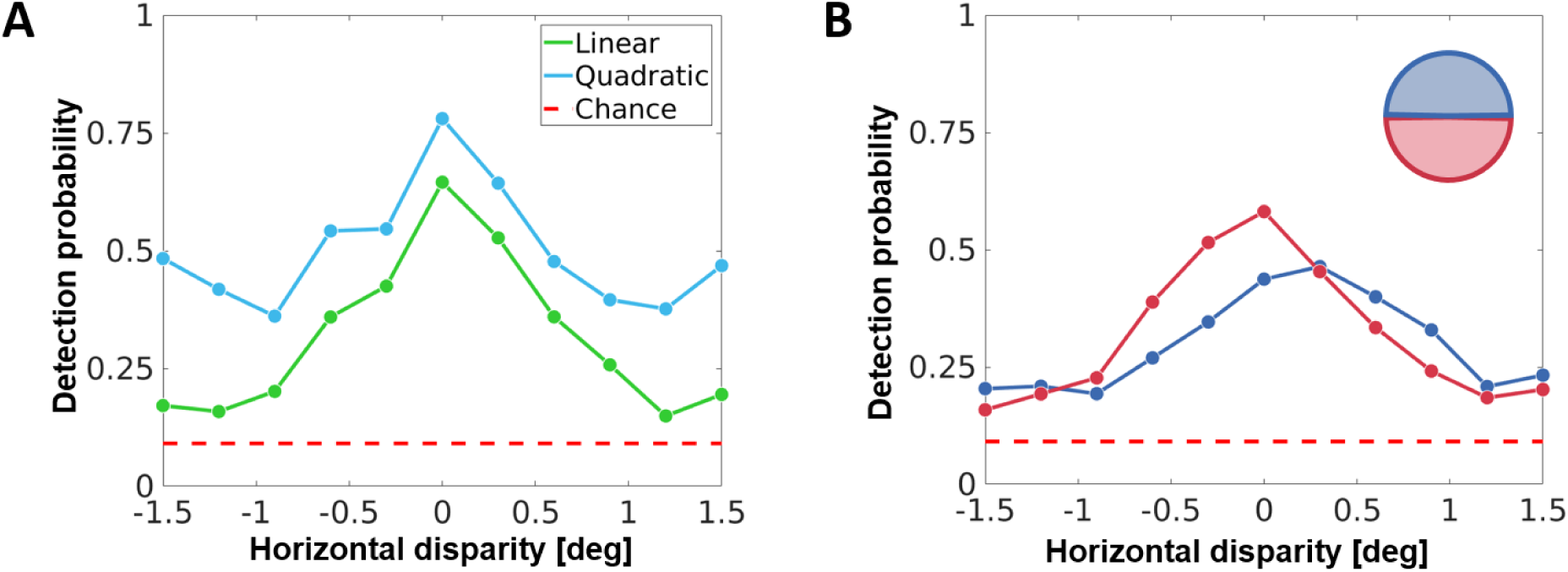
Decoding disparity through random-dot stereograms (RDS). **A**. Output spikes of the V1 layer (foveal simulation) to RDS stimuli at various disparities were used to train linear (green) and quadratic (blue) discriminant classifiers. The RDS stimuli were chosen as they do not contain any other information except disparity. The performance of these classifiers was then tested using different sets of labelled RDS stimuli (10000 total RDS stimuli; 70% used for training, 30% used for testing; cross-validation using 25 folds). This plot shows the detection probability (probability of correct identification), plotted against the disparity. The quadratic classifier performs better than the linear classifier, while both classifiers perform above chance-level (in dashed red). **B**. Two populations of V1 neurones trained on the upper (blue) and lower (red) hemi-fields were used to test a linear classifier using the same stimuli as **A**. The detection probability of the two classifiers reflects the population bias induced by the disparity statistics of the upper and lower hemifields (Figure 5B).

We have previously seen that V1 populations trained on the upper and lower hemi-fields show systematic biases for uncrossed and crossed disparities respectively (see Figure 5B). In Figure 7B, we show the performance of two linear classifiers, trained on V1 populations from the upper (blue) and lower (red) hemi-fields using the same method as Figure 7A. The crossed/uncrossed disparity biases in the V1 layer are also transferred to the decoding performance of the classifiers. Thus, our model predicts that biases in early binocular neural populations can drive downstream disparity discrimination biases too – something which has indeed been reported in human subjects (Sprague et al., 2015).

## 4. Discussion

We have demonstrated that unsupervised Hebbian learning from natural stereoscopic scenes leads to a population of neurones which are capable of encoding binocular disparity statistics in natural scenes. Our findings provide evidence that known mechanisms of neuronal Hebbian plasticity (such as STDP) could play a key role in the encoding of the statistics of natural scenes in the early visual pathway.

Starting from random receptive-fields, our neural population converges to Gabor-like receptive fields found in V1 simple cells (see Movie 1 and Figure 2). This convergence is achieved by a Hebbian pruning of the initial densely connected dendritic receptive field of size 3° × 3°. The mean size of the converged binocular receptive fields when described using a Gabor fit is about 1° (average 1*σ* of the Gaussian envelope). Although in the reported foveal population we started with a 3° × 3° receptive field, simulations with an initial receptive field size of 6° × 6° yielded very similar results. Thus, the pruning process is independent of the size of the initial dendritic tree, and yields converged receptive fields which correspond to about 3 times the size of the central ON/OFF region of the retinal ganglion receptive fields.

We find that the converged receptive fields in the two eyes show strong correlations for properties such as the orientation, size and frequency (Figure 3C shows the orientation, but similar results were found for size and frequency). This suggests that the selectivity of the neurones is driven by the inter-eye correlations, despite competition from within-eye correlations. Although correlation-based refinement of binocular receptive-fields has previously been proposed at a theoretical level (Berns et al., 1993), our results prove that natural scenes and the geometry of the human visual system are indeed capable of providing the strength of inter-ocular correlation required for binocularity to occur. This point is further illustrated by our simulation of the oculomotor behaviour in strabismic amblyopes where the inputs are binocular, but the inter-ocular correlations are weakened due to uncoordinated eye fixations. Whilst this population of neurones also converges to Gabor-like receptive fields, a majority of the neurones are found to be monocular (Figure 3B). This is not altogether surprising as STDP learning from monocular images is known to produce oriented Gabor-like receptive fields (Delorme et al., 2001; Masquelier, 2012). These results also echo findings in monkeys reared with amblyopic deficits (Movshon et al., 1987; Kiorpes et al., 1998) where the binocularity of V1 neurones is found to be severely restricted.

The converged neurones show selectivity to zero, near and far disparities between −0.5° and 0.5° (Figure 3D shows this for position disparity, and a similar range is observed when relative phase values are converted to visual angles), which is in line with electrophysiological findings (Poggio et al., 1985; Prince et al., 2002a), and gives the population the theoretical capacity to encode a large range of realistic disparity values (Sprague et al., 2015). Through retinotopic simulations, we give two further examples which show that when allowed to learn from specific regions of the visual field, our model produces population biases which correspond closely to retinotopic biases reported in electrophysiology. In the first example (Figure 5A), we demonstrate that neurones trained on stimuli from peripheral eccentricities are selective to a larger range of horizontal disparities (Durand et al., 2002) when compared with neurones trained on foveal stimuli. In the second example (Figure 5B) we use two neural populations (upper and lower hemi-field) to show how Hebbian learning can lead to neural populations which reflect the intuitive bias that objects in the lower hemi-field are more likely to be closer than fixation and objects in the upper hemifield are likely to be further away. This bias has been demonstrated through free-viewing tasks, and is also reflected in neural responses at both the single-unit (Sprague et al., 2015) and macroscopic (Nasr and Tootell, 2016) levels. Through these simulations, we show that retinotopic learning based on a Hebbian approach could explain how biases in natural statistics can transfer to neural populations in the early visual system.

Few studies have previously characterised the relationship between natural statistics and binocular disparity responses in the visual cortex. Based on their approach to learning, they can roughly be classified under two main headings – unsupervised strategies (Hoyer and Hyvärinen, 2000; Hunter and Hibbard, 2015) which learn from unlabelled stimuli, and supervised approaches (Burge and Geisler, 2014; Goncalves and Welchman, 2017) which use labelled data to learn specific properties of the scene. Whilst results from unsupervised learning can be interpreted as possible encoding schemes for the stimuli, supervised approaches optimise model performance on specific tasks such as disparity or contrast discrimination. Both these approaches solve an optimisation problem which may involve global properties like entropy and mutual information, or task-specific metrics such as classification accuracy. Although our model does not require supervision, it differs critically from traditional, objective-function based unsupervised approaches as it does not necessarily converge towards a static global optimum - instead, it can be interpreted as a self-regulating coincidence detector based on causal Hebbian updates. In our case, the convergence towards an optimum is a result of the stability of natural statistics across samples.

A commonly employed unsupervised approach to studying the encoding of 3D natural statistics is the Independent Component Analysis (ICA) (Hoyer and Hyvärinen, 2000; Hunter and Hibbard, 2015), a technique which maximises the mutual information between the stimuli and the activity of the encoding units. Here, we mention three critical differences which separate our results from those obtained using ICA. First, Hebbian learning could potentially be sub-optimal when viewed in terms of the mutual-information based objective functions used for ICA. For example, Hebbian learning leads to blobby receptive fields which are highly feature invariant, and a very inefficient choice for encoding natural images. Second, Hebbian learning is usually unable to converge to features which do not recur in the input, whereas efficient coding approaches learn the optimal basis representations of the data which could be very different from any individual stimulus. As an example, the converged population in our model contains very few neurones with antiphase receptive fields in the two eyes (Figure 3D). This is in line with electrophysiological data collected in monkeys, cats and the barn owl (Prince et al., 2002a). This result is also very intuitive because in everyday experience, the probability of a given object projecting antiphase features in the two eyes is very low. In contrast, ICA models usually predict a higher number of antiphase receptive fields in the neural population (Hunter and Hibbard, 2015). Third, the receptive fields obtained through the Hebbian model are closer in structure to electrophysiological data than those obtained by ICA (Figure 4) – for example, they have fewer lobes and show broader orientation tuning. Thus, although ICA representations offer a highly optimal encoding of the data in terms of information transfer, they are inconsistent with electrophysiological findings which often include neurones which are sub-optimal in terms of any particular coding strategy (Ringach, 2002).

As initially pointed out by Barlow (1961), although encoding schemes offer explanations as to how scene statistics may be represented in early processing, the precise nature of the encoding must be capable of driving upstream processes which ultimately determine behaviour. We demonstrate (Figure 7) that the activity of the converged V1 population in our model is capable of driving simple classifiers to an above chance performance. This suggests that disparity encoding in the early visual system does not necessarily need supervised training, and an unsupervised feedforward Hebbian process can lead to neural units whose responses can be interpreted in terms of 3D percept through downstream processing. It must be noted that here, decoding through a classifier is only presented as a simplified illustration of perceptual responses, which may involve other processes such as cortico-cortical interactions and feedback.

We have shown that a simple Hebbian learning rule expressed through STDP could provide a plausible mechanism for the encoding of natural statistics in the early visual system. This form of learning through coincidence detection could present a more intuitive explanation for the emergence of representations found in the early visual system and address crucial discrepancies between efficiency based approaches and electrophysiological data (Ringach, 2002). Although this study demonstrates the relevance of Hebbian plasticity in binocular vision, this approach could easily be extended to other sensory modalities, and even multi-modal representations in the early sensory cortex. Finally, we would like to point out that although the present work focuses on the encoding properties of model, this simple plasticity rule also makes it possible to study the time-course of changes in the connectivity of individual units/neurones. For computational neuroscientists, this offers a critical advantage over conventional optimisation-based approaches as it allows for models which track cortical connectivity over time – such as studies on development, critical-period plasticity, and perceptual learning.

## Acknowledgements

We thank Simon J. Thorpe and Lionel Nowak for their comments on the study. This research was supported by the IDEX Emergence (ANR-11-IDEX-0002-02, BIS-VISION) and ANR JCJC (ANR-16-CE37-0002-01, 3D3M) grants awarded to B.R.C., and the FRM grant (FRM: SPF20170938752) awarded to T.C.

